# Investigating the effect of Target of Rapamycin kinase inhibition on the *Chlamydomonas reinhardtii* phosphoproteome: from known homologs to new targets

**DOI:** 10.1101/310102

**Authors:** Emily G. Werth, Evan W. McConnell, Inmaculada Couso, Zoee Perrine, Jose L. Crespo, James G. Umen, Leslie M. Hicks

**Affiliations:** Department of Chemistry, University of North Carolina at Chapel Hill, Chapel Hill, NC; Donald Danforth Plant Science Center, St. Louis, MO 63132, USA; Instituto de Bioquímica Vegetal y Fotosíntesis, Consejo Superior de Investigaciones Científicas (CSIC)—Universidad de Sevilla; Avda. Américo Vespucio, 49, 41092 Sevilla, Spain

**Author notes:** **Correspondence:** Dr. Leslie M. Hicks, Department of Chemistry, University of North Carolina at Chapel Hill, 125 South Road, CB#3290, Chapel Hill, NC 27599, /919-962-2388.

**Keywords:** Phosphoproteomics, Chlamydomonas, AZD8055, rapamycin, Torin1, target of rapamycin, TOR, NL

## Abstract

- Target of Rapamycin (TOR) kinase is a conserved regulator of cell growth whose activity is modulated in response to nutrients, energy and stress. Key proteins involved in the pathway are conserved in the model photosynthetic microalga *Chlamydomonas reinhardtii*, but the substrates of TOR kinase and downstream signaling network have not been elucidated. Our study provides a new resource for investigating the phosphorylation networks governed by the TOR kinase pathway in Chlamydomonas.
- We used quantitative phosphoproteomics to investigate the effects of inhibiting Chlamydomonas TOR kinase on dynamic protein phosphorylation. Wild-type and AZD-insensitive Chlamydomonas strains were treated with TOR-specific chemical inhibitors (rapamycin, AZD8055 and Torin1), after which differentially affected phosphosites were identified.
- Our quantitative phosphoproteomic dataset comprised 2,547 unique phosphosites from 1,432 different proteins. Inhibition of TOR kinase caused significant quantitative changes in phosphorylation at 258 phosphosites, from 219 unique phosphopeptides.
- Our results include Chlamydomonas homologs of TOR signaling-related proteins, including a site on RPS6 with a decrease in phosphorylation. Additionally, phosphosites on proteins involved in translation and carotenoid biosynthesis were identified. Follow-up experiments guided by these phosphoproteomic findings in lycopene beta/epsilon cyclase showed that carotenoid levels are affected by TORC1 inhibition and carotenoid production is under TOR control in algae.

## Introduction

The Target of Rapamycin (TOR) protein kinase is a conserved eukaryotic growth regulator whose activity is modulated in response to stress, nutrients and energy supply (Wullschleger *et al*., 2006; Loewith & Hall, 2011; Dobrenel *et al*., 2016a; González & Hall, 2017; Pérez-Pérez *et al*., 2017). In metazoans and fungi, TOR is found in two compositionally and functionally distinct multiprotein complexes (TORC1) and (TORC2) that control rates of biosynthetic growth and cytoskeletal dynamics respectively (Raught *et al*., 2001; Wullschleger *et al*., 2006). In the green lineage (algae and land plants), only homologs of TORC1 proteins have been identified (Diaz-Troya *et al*., 2008; van Dam *et al*., 2011; Dobrenel *et al*., 2016a). TORC1 kinase activity is modulated by nutrients and stress, and serves to control protein biosynthesis and other metabolic processes in response to environmental conditions (Raught *et al*., 2001). Selective chemical inhibitors of TOR kinase including rapamycin, AZD8055, and Torin1 have been instrumental in dissecting the TOR signaling pathway (Fingar & Blenis, 2004; Thoreen *et al*., 2009; Chresta *et al*., 2010; Benjamin *et al*., 2011). Rapamycin (Rap) inhibits TORC1 activity through an allosteric mechanism requiring formation of a FKBP12-Rap complex (Heitman *et al*., 1991; Brown *et al*., 1994; Sabatini *et al*., 1994). Recent studies support the notion that several functions of TOR kinase are not inhibited by rapamycin (Thoreen *et al*., 2009). Instead, novel drugs like Torin1 and AZD8055 have been reported to more completely inhibit TOR kinase by acting as ATP-competitors (Thoreen *et al*., 2009; Chresta *et al*., 2010). Torin1 has slower off-binding kinetics than other mTOR inhibitors in mammalian cell lines, possibly due to conformational change induction in the kinase that is energetically more difficult to recover from leading to a more pronounced and longer inhibition of the TORC1 pathway (Liu *et al*., 2013). AZD8055 is an ATP-competitive inhibitor of mTOR and all PI3K class I isoforms noted to inhibit the mTORC1 and mTORC2 substrate phosphorylation (Roohi & Hojjat-Farsangi, 2017). These drugs were used to inhibit TOR activity in plants where rapamycin treatment is not highly effective (Zhang *et al*., 2011; Montane & Menand, 2013).

The role of TOR in mammalian and fungal cell metabolism has been extensively investigated (Wullschleger *et al*., 2006; Dibble & Manning, 2013; Saxton & Sabatini, 2017), while its role in photosynthetic eukaryotes is less well established (Zhang *et al*., 2013; Xiong & Sheen, 2014; Dobrenel *et al*., 2016a). TOR has been shown to control growth, metabolism and life span in the model plant *Arabidopsis thaliana* (Arabidopsis) (Dobrenel *et al*., 2011; Ren *et al*., 2012; Xiong, Y. & Sheen, J., 2012; Xiong *et al*., 2013) where the TOR gene is essential (Menand *et al*., 2002). The model green alga *Chlamydomonas reinhardtii* (Chlamydomonas) has key TORC1 complex proteins encoded by single-copy genes including TOR (Cre09.g400553.t1.1), regulatory associate protein target of rapamycin (RAPTOR) (Cre08.g371957.t1.1), and lethal with sec-13 protein 8 (LST8) (Cre17.g713900.t1.2) (Diaz-Troya *et al*., 2008; van Dam *et al*., 2011). Treatment of Chlamydomonas cultures with rapamycin has been shown to slow but not completely arrest cell growth (Crespo *et al*., 2005), activate autophagy (Perez-Perez *et al*., 2010), and induce lipid droplet formation (Imamura *et al*., 2015; Rodrigues *et al*., 2015). Recent work reported a connection between TOR kinase and inositol polyphosphate signaling that governs carbon metabolism and lipid accumulation (Couso *et al*., 2016). Chlamydomonas cells are sensitive to Torin1 and AZD8055 that are potent inhibitors of cell growth at saturating doses (Couso *et al*., 2016) and induce triacylglycerol accumulation (Imamura *et al*., 2016). However, the TOR pathway in Chlamydomonas has yet to be extensively characterized and, to date, only a limited number of candidate TOR kinase substrates have been identified.

We characterized the phosphoproteome of Chlamydomonas that produced a conservative estimate of 4,588 phosphoproteins / 15,862 unique phosphosites (Wang *et al*., 2014) through a qualitative strategy involving extensive fractionation and complementary enrichment strategies, and have now developed label-free quantification (LFQ) to allow simultaneous quantification of 2,547 Chlamydomonas phosphosites (Werth *et al*., 2017). Herein we characterized the effects of TOR inhibition on the Chlamydomonas phosphoproteome. Cultures treated with saturating doses of different TOR inhibitors (rapamycin, AZD8055 and Torin1) revealed hundreds of affected phosphosites with a significant overlap observed between those seen with different inhibitors. Phosphosites from an AZD-resistant mutant were compared with wild type after AZD treatment revealing very few potential off target effects. Hierarchical clustering was used to classify sites and motif analysis was used to assess consensus motifs in clusters.

## Materials and Methods

### Cell culturing and drug treatment

Strain CC-1690 wild-type mt+ (Sager 21 gr) (Sager, 1955) was used for the wild-type Chlamydomonas analysis across all chemical inhibitors. For the control AZD-insensitive strain experiments, strain was obtained from the Umen laboratory (Donald Danforth Plant Science Center). All cultures were maintained on TAP (Tris acetate phosphate) agar plates and grown in 350-mL TAP liquid cultures at 25°C as previously described (Couso *et al*., 2016). Experiments were done using five replicate cultures grown to exponential phase (1-2x10^6^ cells/mL) for each drug condition and control and quenched with 40% methanol prior to harvesting by centrifuging at 4000 *g* for 5 min and discarding supernatant. To limit batch effects, replicate “n” of each drug and control were harvested together (Figure 1) prior to downstream processing. Cell pellets were then flash frozen using liquid nitrogen and stored at -80°C until use. For AZD8055-, Torin 1-, and rapamycin-treated (LC Laboratories) cultures, drug was added to a final concentration of 500 nM for rapamycin and Torin 1, and 700 nM for AZD8055 from 1mM stocks in DMSO for 15 min prior to harvesting. For control replicates, just drug vehicle (DMSO) without a chemical inhibitor was added to each replicate culture for 15 min prior to harvesting.

**Figure 1.**
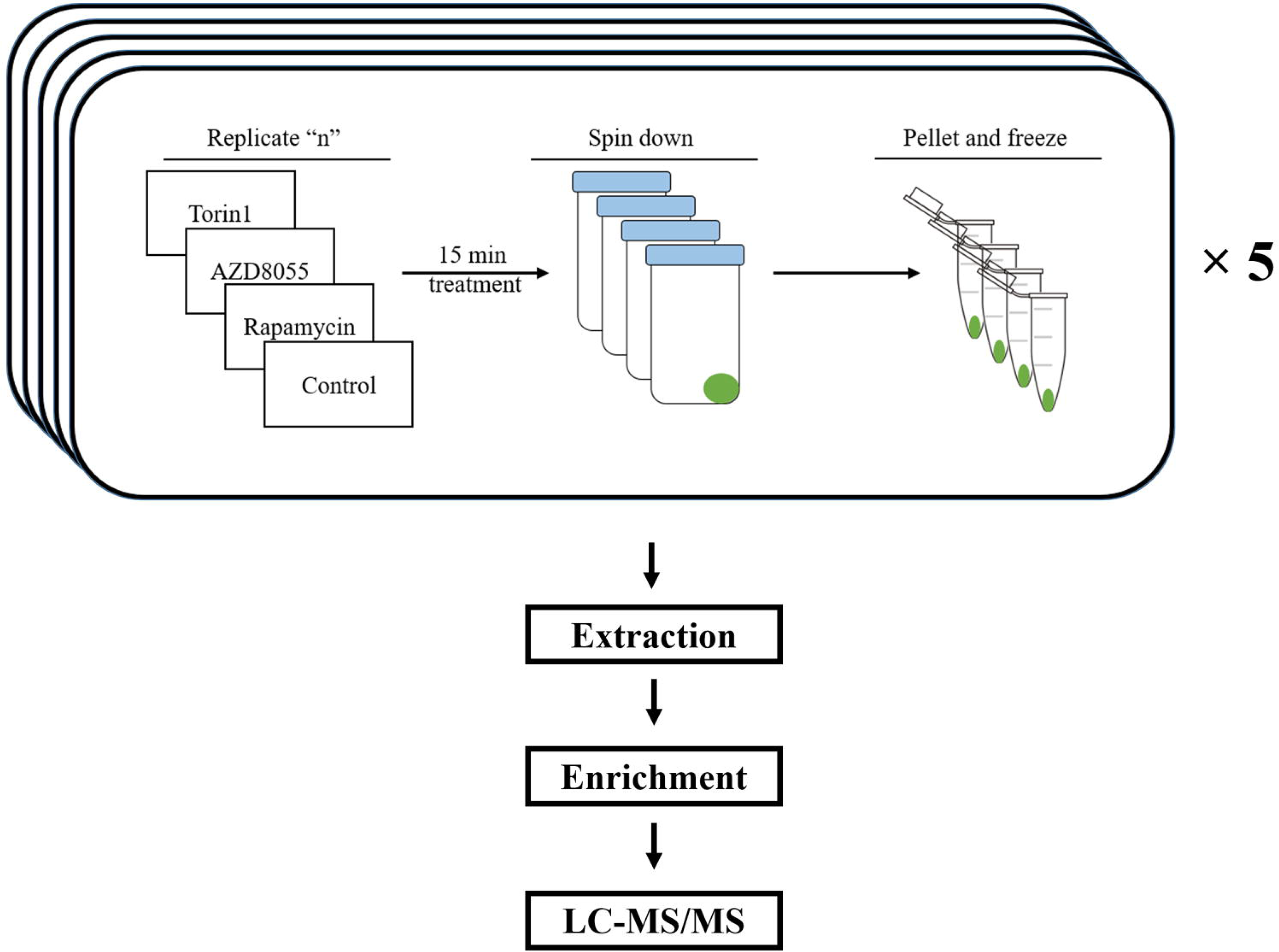
Drug treatment and cell harvesting workflow in Chlamydomonas cells. Replicate “n” (1-5) of each drug condition and control were harvested together prior to downstream processing. To minimize inter-condition batch effects, “n” replicate of each condition was harvested together and frozen until protein extraction.

### Protein extraction

Cell pellets were resuspended in lysis buffer containing 100 mM Tris, pH 8.0 with 1x concentrations of cOmplete protease inhibitor and phosSTOP phosphatase inhibitor cocktails (Roche, Indianapolis, IN, USA). Cells were lysed via sonication using an E220 focused ultrasonicator (Covaris, Woburn, MA, USA) for 120 s at 200 cycles/burst, 100 W power and 13% duty cycle. Following ultrasonication, the supernatant was collected from cellular debris by centrifugation for 10 min at 15,000 *g* at 4°C and proteins were precipitated using 5 volumes of cold 100 mM ammonium acetate in methanol. Following 3 hr incubation at -80°C, protein was pelleted by centrifugation for 5 min at 2,000 *g* followed by two washes with fresh 100 mM ammonium acetate in methanol and a final wash with 70% ethanol. Cell pellets were resuspended in 8M urea and protein concentration was determined using the CB-X assay (G-Biosciences, St. Louis, MO, USA).

### Protein digestion and reduction

Samples were reduced using 10 mM dithiothreitol for 30 min at RT and subsequently alkylated with 40 mM iodoacetamide for 45 min in darkness at RT prior to overnight digestion. Samples were diluted 5-fold in 100 mM Tris following alkylation and digestion was performed at 25C for 16 h with Trypsin Gold (Promega) at a protease:protein ratio of 1:50.

### Solid-phase extraction

After digestion, samples were acidified to pH<3.0 with trifluoroacetic acid (TFA). Pelleted, undigested protein was cleared from the supernatant by centrifugation for 5 min at 5,000 *g* prior to solid-phase extraction. Desalting was performed using C18 50 mg Sep-Pak cartridges (Waters). Columns were prepared by washing with acetonitrile (MeCN) followed by 80% MeCN/20%H2O/0.1% TFA and 0.1% TFA. Digested protein lysates were applied to the columns and reloaded twice before being washed with 0.1% TFA and eluted using 80% MeCN/20%H2O/0.1% TFA.

### Phosphopeptide enrichment and clean-up

Following protein digestion and solid-phase extraction, replicates were dried down using vacuum centrifugation and phosphopeptide enrichment was performed on 2-mg aliquots of each sample using 3 mg Titansphere Phos-TiO2 kit spin columns (GL Sciences) as previously described (Werth *et al*., 2017). After enrichment, samples were dried down and desalted again using ZipTips (Millipore) as per manufacturers protocol prior to LC-MS/MS acquisition.

### LC-MS/MS acquisition and data processing

Following ZipTip clean-up, peptides were dried down and resuspended in 20 μL of 0.1% TFA, 5% MeCN before separation via a 90-min linear gradient from 95% H_2_O/5% MeCN/0.1% formic acid (FA) to 65% H2O/35% MeCN/0.1% FA via a NanoAcquity UPLC (Waters) using a C18 column (NanoAcquity UPLC 1.8 μm HSS T3, 75 μm × 250 mm). A TripleTOF 5600 (AB Sciex) Q-TOF was operated in positive-ionization nanoelectrospray and high-sensitivity mode for data acquisition as previously described (Slade *et al*., 2015). In addition to the Supporting Information tables for MS datasets, the mass spectrometry proteomics data have been deposited to the ProteomeXChange Consortium via PRIDE partner repository(Vizcaíno *et al*., 2013) identifier PXD007221. Acquired spectra (*.wiff) files were imported into Progenesis QI for proteomics (v2.0, Nonlinear Dynamics) as previously described (Werth *et al*., 2017) with peptide sequence determination and protein inference done by Mascot (v.2.5.1; Matrix Science) using the *C. reinhardtii* Phytozome v.11 database (www.phytozome.net/; accessed May 2015) appended with the NCBI chloroplast and mitochondrial databases (19,603 entries) and sequences for common laboratory contaminants (http://thegpm.org/cRAP/; 116 entries). For database searching, trypsin protease specificity with up to two missed cleavages, peptide/fragment mass tolerances of 20 ppm/0.1 Da, a fixed modification of carbamidomethylation at cysteine, and variable modifications of acetylation at the protein N-terminus, oxidation at methionine, deamidation at asparagine or glutamine, phosphorylation at serine or threonine and phosphorylation at tyrosine were used. Peptide false discovery rates (FDR) were adjusted to ≤1% using the Mascot Percolator algorithm (Käll *et al*., 2007) and only peptides with a Mascot ion score over 13 were considered.

Custom scripts written in Python were implemented to parse results following data normalization and quantification in Progenesis QI for proteomics. Shared peptides between proteins were grouped together to satisfy the principle of parsimony and represented in Table S1 by the protein accession with the highest amount of unique peptides, otherwise the largest confidence score assigned by Progenesis QI for proteomics. Additionally, the script appended site localization of variable modifications using an implementation of the Mascot Delta Score (Savitski *et al*., 2011) to the peptide measurements (*.csv) export from Progenesis QI for proteomics with confident site localization considered a Mascot Delta score >90%. Following scoring, only peptides with phosphorylation at serine, threonine, or tyrosine were considered for further processing and analysis.

### Downstream bioinformatics analysis

Missing value imputation was performed on logarithmized normalized abundances in Perseus v1.6.0.0 (Cox & Mann, 2012; Tyanova *et al*., 2016) requiring at least three of the five replicates in all drug conditions and control to be nonzero to continue through the workflow. A coefficient of variation (CV) cutoff was applied requiring CV<25% in at least 2 of 4 conditions for each phosphosite. For t-test analyses, replicates were grouped and the statistical tests were performed with fold change threshold of ±2 and p≤0.05 significance threshold. KEGG pathway annotation (Kanehisa & Goto, 2000), Gene Ontology (GO) (Ashburner *et al*., 2000) term annotation, hierarchical clustering, and motif analysis were performed following statistical testing to glean biological insight on modulated sites found in the study. For hierarchical clustering, visualization was performed in Perseus v1.6.0.0. Following data normalization and missing value imputation, intensity values were z-score normalized and grouped using k-means clustering with default parameters. For motif analysis, sequence logo visualizations were performed using pLOGO with serine or threonine residues fixed at position 0. Positions with significant residue presence are depicted as amino acid letters sized above the red line (O’shea *et al*., 2013).

### Carotenoid analysis

Chlamydomonas cells were collected by centrifugation (4000 g for 5 min) and resuspended in 80% acetone. Samples were heat up for 5 min in a water bath at 90°C and then centrifuge at 10000g 10min. The supernatant evaporated under N2, and then resuspended in 80% acetone. The separation and chromatographic analysis of pigments was performed in a HPLC using a Waters Spherisorb ODS2 column (4.6 x 250 mm, 5μm particle size). The chromatographic method described by Baroli et al., 2003 (Baroli *et al*., 2003). Pigments were eluted at a flow rate of 1.0 mL min^-1^ with a linear gradient from 100% solvent A (acetonitrile:methanol:0.1mM Tris-HCl pH 8.0 [84:2:14]) to 100% solvent B (methanol:ethyl acetate [68:32]) for 20 min, followed by 7 min of solvent B, then 1 min with a linear gradient from 100% solvent B to 100% solvent A, and finally 6 min with solvent A. The carotenoids were detected at 440 nm using a Waters 2996 photodiode-array detector. The different carotenoids were identified using standards from Sigma (USA) and DHI (Germany). This analysis was normalized by dry cell weight. Dry weight was determined by filtering an exact volume of microalgae culture (30 mL) on pre-targeted glass-fiber filters (1μm pore size). The filter was washed with a solution of ammonium formate (0.5 M) to remove salts and dried at 100 °C for 24 h. The dried filters were weighed in an analytical balance and the dry weight calculated by difference.

### SDS-PAGE and Western Blotting

Chlamydomonas cells from liquid cultures were collected by centrifugation (4000 g for 5 min), washed in 50 mM Tris-HCl (pH 7.5), 10 mM NaF, 10 mM NaN3, 10 mM p-nitrophenylphosphate, 10 mM sodium pyrophosphate, and 10 mM b-glycerophosphate), and resuspended in a minimal volume of the same solution supplemented with Protease Inhibitor Cocktail (Sigma). Cells were lysed by two cycles of slow freezing to –80 °C followed by thawing at room temperature. The soluble cell extract was separated from the insoluble fraction by centrifugation (15 000 g for 20 min) in a microcentrifuge at 4 °C. For immunoblot analyses, total protein extracts (20 μg) were subjected to 12% SDS–PAGE and then transferred to PVDF membranes (Millipore). Anti-P-RPS6(Ser242) and anti-RPS6 primary antibodies were generated as described in Dobrenel et al., 2016 (Dobrenel *et al*., 2016b) and produced by Proteogenix, (France). Phospho-p70 S6 kinase (Thr(P)-389) polyclonal antibody (Cell Signaling, 9205) was used as described in Xiong et al., 2012 (Xiong, Yan & Sheen, Jen, 2012). Primary antibodies were diluted 1:2000 and 1:1000 respectively. Secondary anti-rabbit (Sigma) antibodies were diluted 1:5000 and 1:10 000, respectively, in phosphate-buffered saline (PBS) containing 0.1% (v/v) Tween-20 (Applichem) and 5% (w/v) milk powder. The Luminata Crescendo Millipore immunoblotting detection system (Millipore) was used to detect the proteins. Proteins were quantified with the Coomassie dye binding method (BioRad).

## Results

### Parameter selection for TORC1-specific inhibition

Previous studies in Chlamydomonas have shown rapamycin drug saturation ranging from 500 nM-1μM (Crespo *et al*., 2005). For this study, 500 nM rapamycin was selected and saturating doses for Torin1 and AZD8055 in wild-type Chlamydomonas strain CC-1690 were determined using serial dilutions with previously published target concentrations (Couso *et al*., 2016). Growth inhibition saturated at 500 nM for Torin1 and 700 nM for AZD8055 (Supplemental Figure 1).

While reports have shown phosphorylation changes as early as 2 minutes after rapamycin treatment (Rigbolt *et al*., 2014), a 15-minute time point was chosen based on the high number of changes seen in mammalian cell lines at this time point (Demirkan *et al*., 2011; Harder *et al*., 2014; Rigbolt *et al*., 2014) and to ensure reproducibility in treatment and harvesting across 20 samples (control, AZD8055-, Torin1-, and rapamycin-treated with n=5) from the early logarithmic phase of growth. Growth for each replicate was staggered, and to limit batch-effects replicates were harvested in sets, each containing a control sample and the three different drug-tested samples (Figure 1) prior to downstream processing.

Prior rapamycin phosphoproteomic experiments in mammalian studies have shown that phosphopeptide ratios in general were not affected by normalization to protein levels at a 15 min time point (Harder *et al*., 2014). To confirm this in *Chlamydomonas reinhardtii*, a whole-cell proteomics experiment (n=4) was performed after 15 min of rapamycin inhibition. These results showed that protein abundance levels in general are not affected with only 18 of the 1,539 proteins quantified significantly changing (Supplemental Table S4) with no significant differences in protein abundances between control and treatment (Supplemental Figure 2). While 4 of the 18 proteins changing at the protein level were identified in the phosphoproteomics study detailed below, they were not detected as phospho-modulated following chemical inhibition and thus not proteins of interest in this study. Thus, we have confidence that the statistically significant phosphorylation sites detected are from changes in the phosphorylation status and not an artefact of protein expression or turnover.

### Quantitative coverage of the TOR-inhibited phosphoproteome

Label-free quantitative phosphoproteomics was used to compare normalized abundance values of control samples (n=5) versus samples treated with each of the chemical inhibitors (n=5) using an area under the curve (AUC) MS1 intensity-based quantitation method. For this approach, the change in chromatographic peak area between control and chemically-inhibited replicates for each phosphopeptide was the basis for determining relative phosphopeptide abundance. Tip-based TiO2 phosphopeptide enrichment that previously showed high reproducibility between samples (Werth *et al*., 2017) was used for sample preparation. As part of the LFQ pipeline, quantitative data was filtered for only peptides containing a phosphorylation site on Ser, Thr, or Tyr after peak picking and peptide sequence determination. At least 3 of the 5 replicates for each condition were required to have nonzero abundances to remain in the final dataset presented in Table S1 and missing value imputation was performed on log-transformed normalized abundances (Cox & Mann, 2012; Tyanova *et al*., 2016). Highly variable sites remaining in the dataset were then removed by filtering out those with a coefficient of variation of >25% in >2 experimental conditions. The resulting dataset contained 2,547 unique phosphosites from 1,432 different proteins (Table S1) in untreated control samples. To determine sites of interest following chemical inhibition with Torin1, AZD8055, or rapamycin, two sample Student’s T-tests were performed between samples from each chemical inhibitor compared and control samples. From this, 258 phosphosites from 219 phosphopeptides showed at least a two-fold change and a p-value ≤ 0.05 (Figure 2a, Table S2). High confidence phosphorylation site assignments (90% site-localization based on Mascot Delta scoring(Savitski *et al*., 2011)) were achieved for 48% of the dataset (1,123 of the 2,363 phosphopeptides) listed in Table S1. AZD8055 treatment resulted in 97 phosphopeptides modulated in the wild-type strain (Figure 2a). A matched control experiment using an AZD-insensitive strain which grow similar to wild-type (Supplemental Figure 3) showed only 13 low abundance phosphosites differentially changing (Table S3, Figure 2b). Of the 13, no overlap was found with the 258 modulated phosphosites in the main dataset.

**Figure 2:**
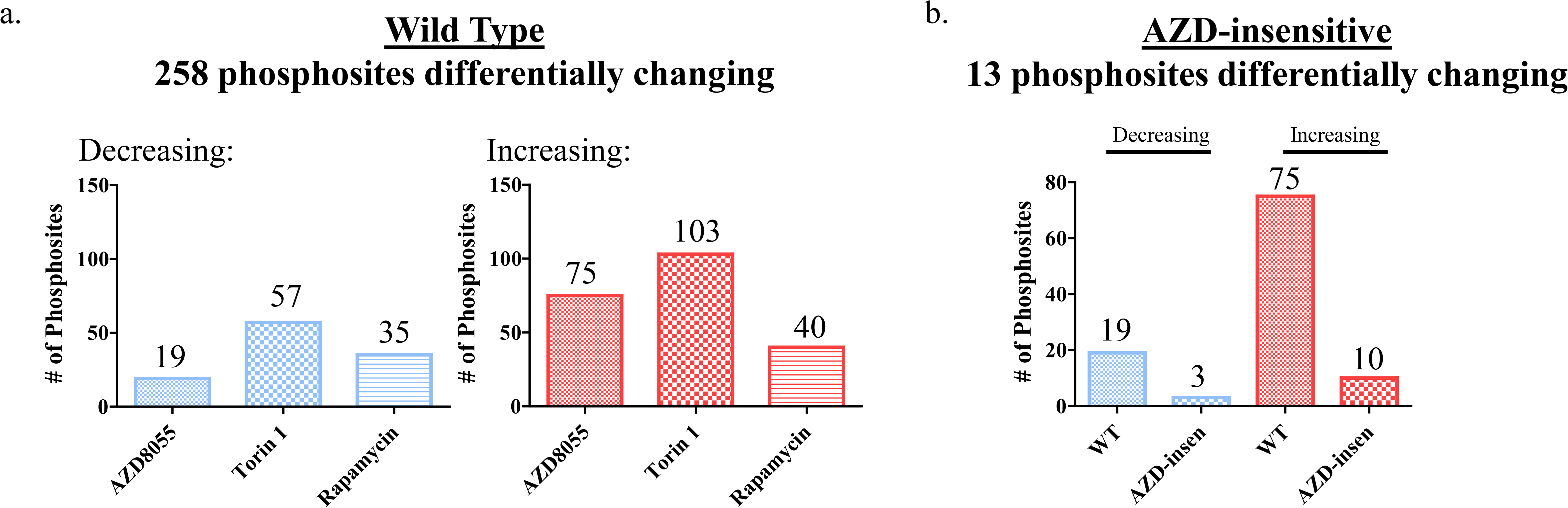
Sites modulated by TOR inhibition. Results of differential analysis between each chemical inhibitor drug treatment compared to control for both wild-type (a) and AZD-insensitive (b) Chlamydomonas strains. For comparison of overlap between the drug conditions in the WT dataset, a Pearson’s correlation was performed comparing all condition types. From this, the highest correlation among conditions was between AZD8055 and Torin1 at 0.986 and the lowest 3 were all drug inhibitor vs. controls.

Torin1 treatment caused the largest number of significant changes with 103 up- and 57 down-modulated phosphosites. AZD8055 treatment caused 75 up- and 19 down-modulated phosphosites, while rapamycin treatment caused 40 up- and 35 down-modulated phosphosites. Overlap analysis of the differential sites for each drug revealed 88% (57/66) of all the down-modulated sites were in the Torin1 subset, while 42% (24/57) of the Torin1 down-modulated sites were not detected with AZD or rapamycin. Up-regulated sites were also compared for each condition and to determine if the conditions had significant overlap between down- and up-modulated sites, a hypergeometric test was performed with p-values of 3.76x10^-25^ and 2.87 x10^-34^, respectively, showing significant overlap.

### Cluster analysis and phosphosite motif identification

Kinase specificity can be dictated by amino acid residues immediately surrounding phosphorylation sites on substrates (Chou & Schwartz, 2011). Mammalian TOR has been shown to mainly (but not exclusively) phosphorylate (S/T)P motifs and motifs with hydrophobic residues surrounding the phosphorylation site making it a relatively promiscuous kinase whose substrate choices may also be influenced by additional interactions outside the phosphosite region (Robitaille *et al*., 2013). Hierarchical clustering of Chlamydomonas modulated phosphosites generated 2 distinct clusters (Figure 3a,b), and motif analysis (O’shea *et al*., 2013) was performed on decreasing (cluster 1) and increasing (cluster 2) clusters. Cluster 1 phosphosites, which contained 94% of sites that significantly decrease in phosphorylation upon TOR inhibition, had significant enrichment for a proline in the +1 position and arginine in the -3 position with respect to the phosphorylation site (position 0) that showed strong enrichment for serine over threonine (Figure 3c). Cluster 2 phosphosites also had significant enrichment for a proline in the +1 position and arginine in the -3 position in addition to enrichment for an aspartic acid at the +3 position. Thus, CrTOR may have a preference for phosphorylation of (S/T)P motifs on substrates, similar to mTOR(Robitaille *et al*., 2013) and other diverse proline-directed kinases including cyclin-dependent protein kinases (CDKs) and mitogen-activated protein kinases (MAPKs) (Lu *et al*., 2002). Additionally, a phosphoproteomic study using mammalian cell line MCF7 identified the RXXS/TP motif identified in clusters 1 and 2 as a rapamycin-sensitive motif (Rigbolt *et al*., 2014). Other studies have also found RXRXXS/T and RXXS/T motifs (Demirkan *et al*., 2011; Harder *et al*., 2014) enriched among rapamycin-sensitive phosphosites that are recognized by mTOR-regulated kinases Akt, S6K1 and SGK1 (Hsu *et al*., 2011). Cluster 2 additionally has an acidic motif also found in casein kinase-II substrates (Lv *et al*., 2014).

**Figure 3.**
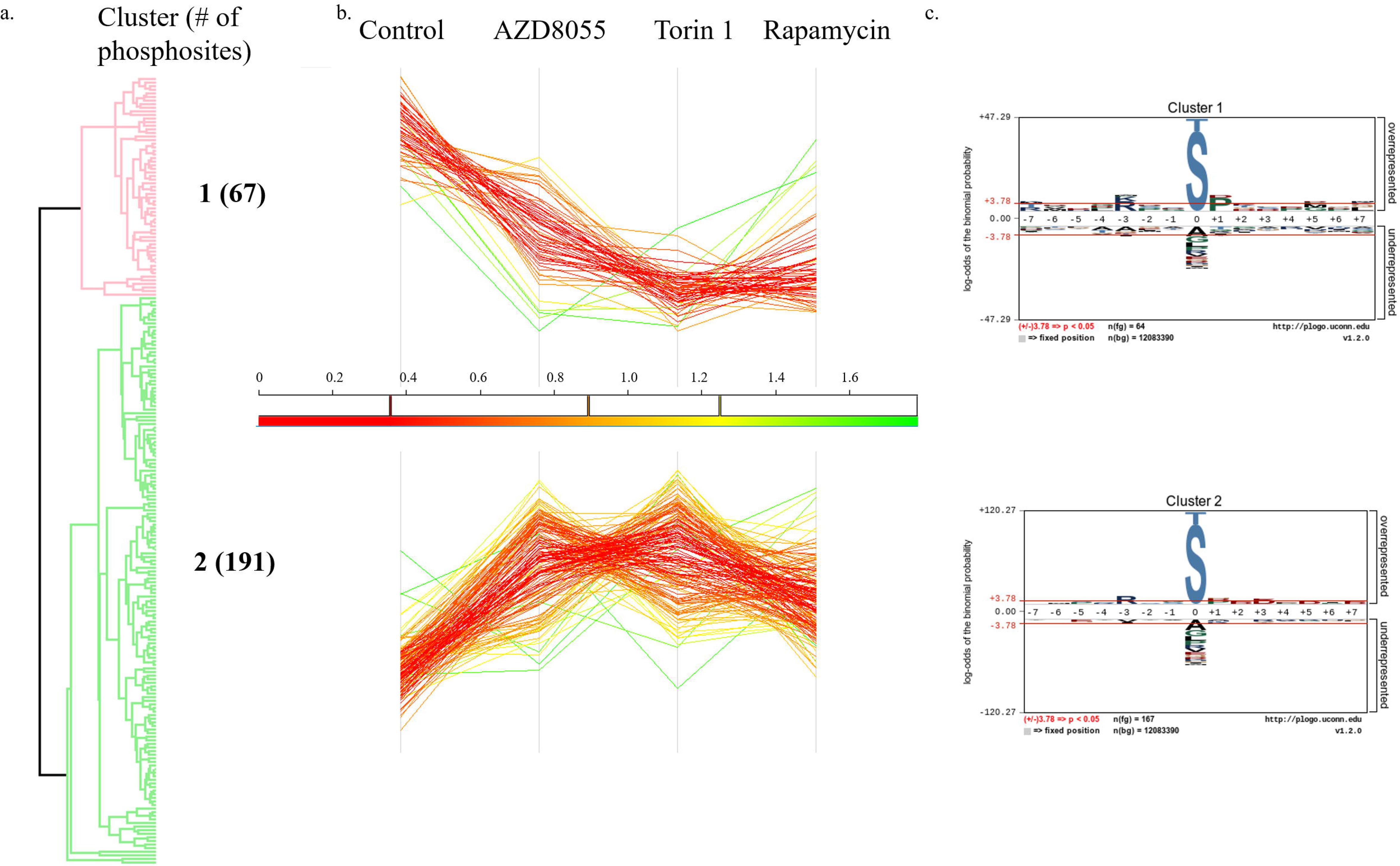
Hierarchical clustering of differentially changing sites into 2 clusters (a). Visualization was performed in Perseus v1.6.0.0. Following data normalization and missing value imputation, intensity values were z-score normalized and grouped using k-means clustering with default parameters. Overall trends in site intensity were graphed and colored based on intensity (b). For each of the two clusters, motif analysis was performed (c). Sequence logo visualizations were performed using pLOGO with serine or threonine residues fixed at position 0. Positions with significant residue presence are depicted as amino acid letters sized above the red line. For cluster 1, there was significant enrichment for a proline in the +1 position and arginine in the -3 position, RXXS/TP. For cluster 2, there was again significant enrichment for a proline in the +1 position and arginine in the -3 position in addition to an aspartic acid in the +3 position, RXXS/TPXD.

### Phosphosites in TORC1 complex proteins

Numerous phosphosites in mammalian homologs of TORC1 complex proteins are regulated by the TOR pathway and/or are phosphorylated autocatalytically (Foster *et al*., 2010). This includes sites on Raptor and mTOR homologs. Therefore, phosphosites found on CrTORC1 complex proteins could be affected by TOR inhibition. TORC1 complex proteins conserved in Chlamydomonas include TOR (Cre09.g400553.t1.1), Raptor (Cre08.g371957.t1.1), and LST8 (Cre17.g713900.t1.2) (Merchant *et al*., 2007; Diaz-Troya *et al*., 2008; Perez-Perez *et al*., 2010; Couso *et al*., 2016). While there is a known LST8 homolog in Chlamydomonas, it is not known to be phosphorylated (Wang *et al*., 2014). Phosphosites on Raptor (Ser782/783:NL) (Not Localized:NL) and TOR (Ser2598) were detected in this study, however no statistically significant modulation in their abundance was detected. BLASTP alignment of human Raptor (Uniprot Q8N122) with CrRaptor revealed high sequence overlap on the N-terminal region of the protein (residues 9-627 with 57% identity), however known TORC1-sensitive phosphosites in the human Raptor homolog (i.e. Ser719, Ser721, Ser722, Ser859, and Ser863 (Carrière *et al*., 2008; Foster *et al*., 2010)) were not conserved in CrRaptor. Similarly, human mTOR (Uniprot P42345) phosphosites Ser2159/Thr2164 that are within the kinase domain promoting mTORC1-associated mTOR Ser2481 autophosphorylation (Ekim *et al*., 2011) are not conserved in CrTOR. The limited sequence conservation among CrTORC1 phosphosites with mammalian TOR phosphosites precludes any predictions about functions of CrTORC1 protein phosphorylation. Other phosphosites on CrTORC1 complex proteins that were detected in previous work on the global phosphoproteome in Chlamydomonas (Wang *et al*., 2014) might be significant for regulation but they were not observed in our data. Future experiments with additional fractionation to increase the dynamic range of quantitative coverage could allow for deeper coverage and more comprehensive detection of phosphosites.

## Discussion

### Sites modulated by TORC1 inhibition – known and putative substrates

In animal cells TORC1-inhibition blocks phosphorylation of multiple substrates including S6 kinases and eukaryotic translation initiation factors, leading to a reduction in translation initiation rates for a subset of mRNAs (Jefferies *et al*., 1994; Terada *et al*., 1994; Wang & Proud, 2009). Phosphorylation of Ser371 and Thr389 in human p70S6K1 (Uniprot P23443-2) are reduced by treatment of cells with TOR inhibitors (Dennis *et al*., 1996; Burnett *et al*., 1998). While we identified one potential site (site was not localized) (Thr771/Ser773/Thr777:NL) on a Chlamydomonas homolog of ribosomal protein S6 kinase (S6K; Cre13.g579200.t1.2), its phosphorylation state was not significantly altered by TOR inhibitors (Table 1). No coverage was obtained on predicted conserved sites Ser915 and Thr932, which align to human p70S6K1 Ser371 and Thr389, respectively, although these sites have been detected previously in Chlamydomonas (Wang *et al*., 2014). Moreover, while commercial anti-phospho S6K antibodies have been shown to detect phospho-S6K in plants (Xiong, Yan & Sheen, Jen, 2012; Ahn *et al*., 2014) they have not detected a signal in Chlamydomonas in our hands (Supplemental Figure 4) and in another study (Couso *et al*., 2016), thus limiting our ability to independently validate Chlamydomonas TOR substrate phosphopeptides. On the other hand, Chlamydomonas ribosomal protein S6 (RPS6, Cre09.g400650.t1.2), a predicted target of S6K, showed a 2.1-fold decrease in phosphorylation on Thr127 following Torin1 treatment (Figure 5, Table 1). While this site is potentially TORC1-regulated, antibodies specific for this phosphosite needed for validation are not available. In Arabidopsis, a phosphosite on the C-terminal extremity peptide of RPS6, Ser240, had decreased phosphorylation following TOR inactivation (Dobrenel *et al*., 2016b). While this exact site is not conserved in Chlamydomonas, the phosphoserine next to it, Ser241 in Arabidopsis (aligning to Ser242 in Chlamydomonas) has been detected in prior work (Wang *et al*., 2014); however it was not detected in this study (Figure 4a). To determine if Ser242 in Chlamydomonas is TORC1-regulated, a western blot of proteins fractionated from wild-type cells under different drug treatments for 0, 5, 15, 30, and 60 min was performed with antibodies raised for phosphorylated and non-phosphorylated Ser242 (Figure 4b), the latter used as a control for monitoring protein level. Interestingly, this site does not seem to change drastically with Torin1, AZD8055, or rapamycin treatment contrary to results on the C-terminal phosphosite in Arabidopsis.

**Table 1:**
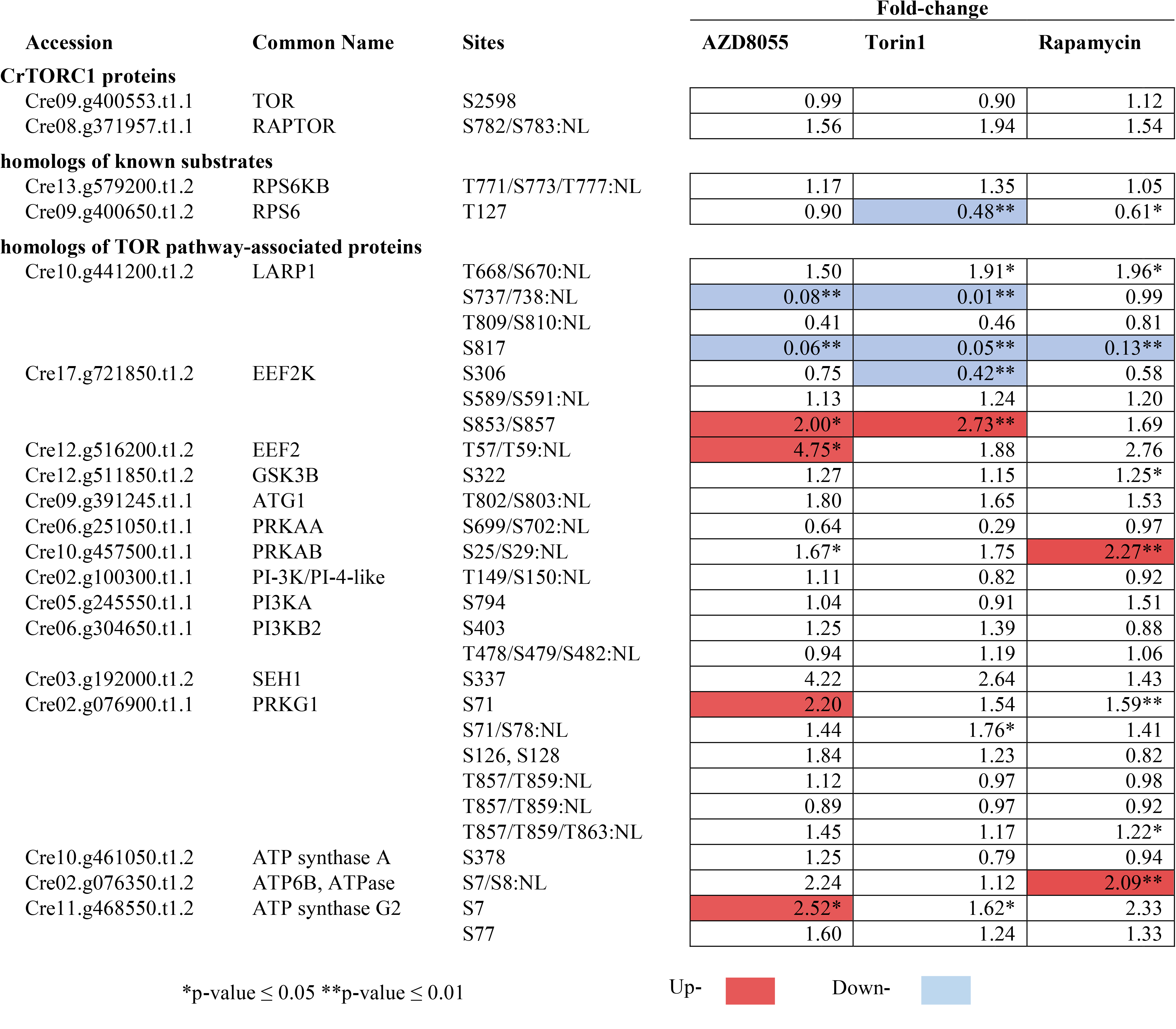
TOR targets identified with fold change values for drug condition versus control. Fold change values shaded red indicate a statistically significant increase in phosphopeptide abundance for specified drug treatment versus control. Fold change values shaded blue indicate a statistically significant decrease in phosphopeptide abudance for specified drug treatment versus control. Level of p-value statistical significance is denoted by p-value ≤ 0.05 (*) and ≤ 0.01 (**)

**Figure 4.**
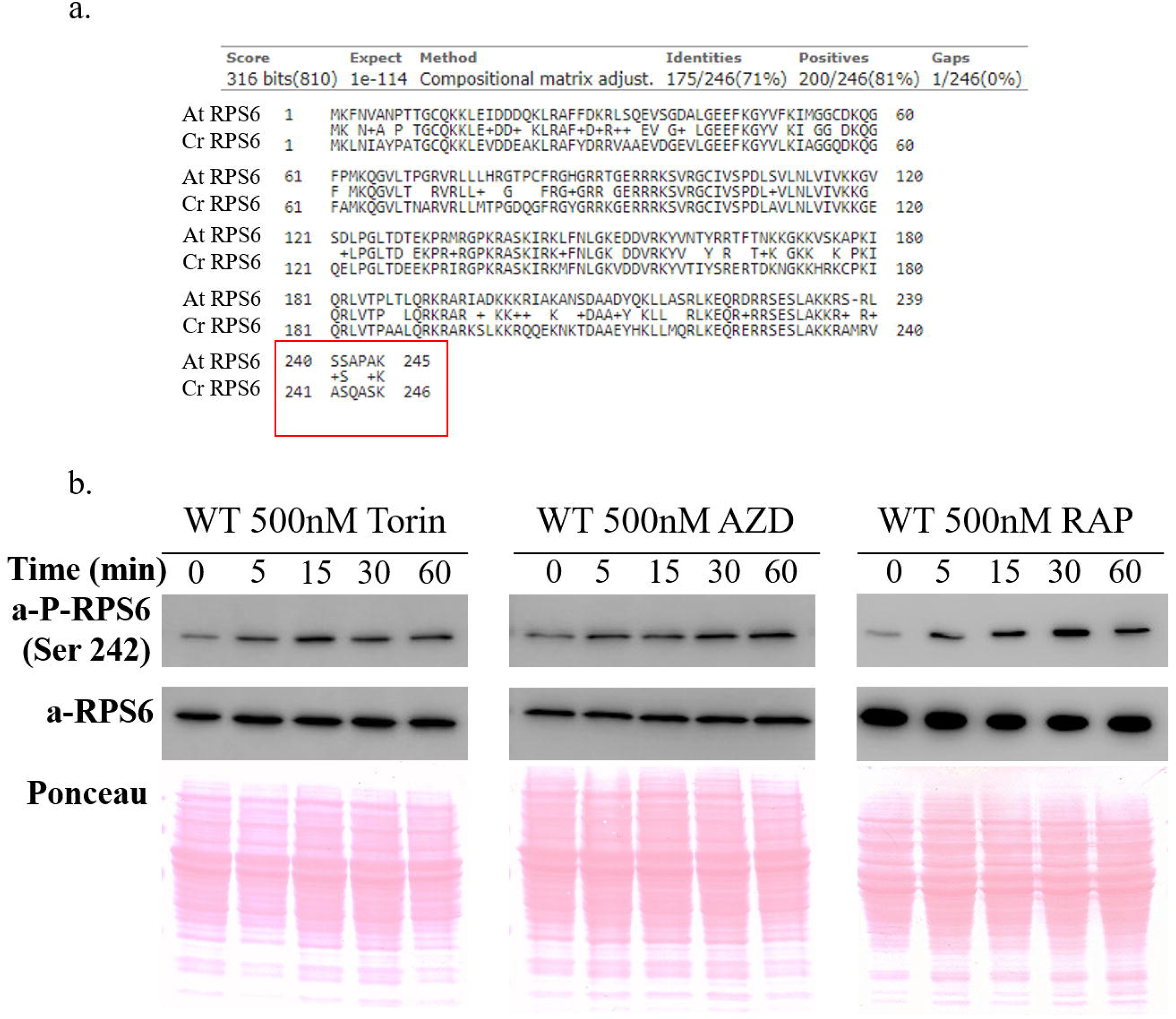
Comparison of RPS6 protein sequence between Arabidopsis and Chlamydomonas (a). a western blot in wild-type under different drug treatments for 0, 5, 15, 30, and 60 min with antibodies raised for Ser242 (b).

### Sites modulated by TORC1 inhibition – known TOR pathway association

Of the 258 phosphosites detected as significantly modulated in this study, 10 are in homologs of proteins associated with the TOR signaling pathway (Figure 5, Table 1). In addition to four sites of decreasing phosphorylation, six proteins related to the TOR pathway had an increase in protein phosphorylation following chemical inhibition. While initially an unexpected observation, similar increases were previously reported for some phosphosites in a phosphoproteomic study of TOR inhibition in mouse liver (Demirkan *et al*., 2011). In our study, sites with increasing phosphorylation after TOR inhibition include elongation factor 2 (EEF2, Cre12.g516200.t1.2) whose animal homologs showed reduced activity upon phosphorylation. In human cells, phosphorylation of EEF2 Thr57 by elongation factor 2 kinase (EEF2K, Cre17.g721850.t1.2) inactivates EEF2 activity, an essential factor for protein synthesis (Hizli *et al*., 2013). This site is conserved in Chlamydomonas EEF2 (Thr57/Thr59:NL) where we detect a 4.75-fold increase in phosphorylation with AZD8055 treatment with a predicted effect of reduced translation initiation rates. From these data we predict that CrTOR signaling may inhibit EEF2 kinase activity, and that this inhibition is relieved in the presence of TOR inhibitors.

**Figure 5:**
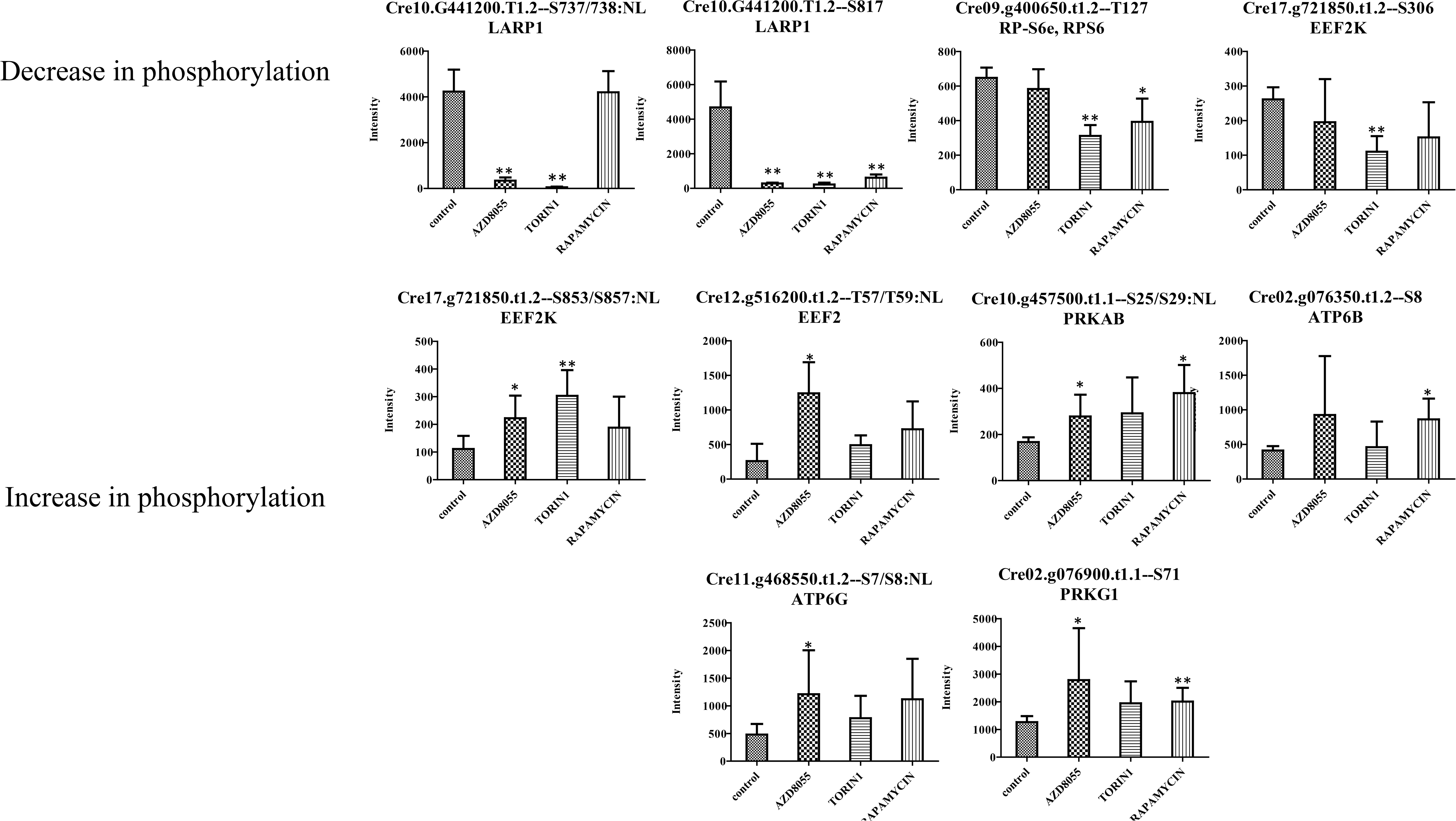
Bar charts of 10 modulated phosphosites on TOR pathway-associated proteins based on homology. Level of p-value statistical significance is denoted by p-value ≤ 0.05 (*) and ≤ 0.01 (**)

LA RNA-binding protein (LARP1, Cre10.g441200.t1.2) had two phosphosites that both underwent large decreases in phosphorylation upon treatment with the three chemical inhibitors. Ser817 was decreased 0.0_6AZD8055_, 0.05Torin1, and 0.13_RAP_ and Ser 737/738:NL was decreased 0.08_AZD8055_ and 0.01_Torin_ 1 but no change in rapamycin (0.99_RAP_) (Figure 5). In mammals, LARP1 phosphorylation also requires mTORC1 (Hsu *et al*., 2011; Yu *et al*., 2011; Kang *et al*., 2013) with studies in human cell lines establishing LARP1 as a target of mTORC1 and S6K with non-phosphorylated LARP1 interacting with both 5’ and 3’ UTRs of RP mRNAs and inhibiting their translation (Hong *et al*., 2017). Additional reports have shown LARP1 as a direct substrate of mTORC1 in mammalian cells with mTORC1 controlling Terminal Oligopyrimidine (TOP) mRNA translation via LARP1 (Fonseca *et al*., 2015; Hong *et al*., 2017). The dramatic modulation of LARP1 phosphorylation detected in our study indicates that LARP1 may have a parallel role in Chlamydomonas. The human LARP1 phosphosites are not conserved with those we found in Chlamydomonas. However, based on the NCBI conserved domain searching (Marchler-Bauer & Bryant, 2004), the DM15 domain required for the interaction of LARP1 with mTORC1 in human cell lines is conserved in Chlamydomonas LARP1, and the phospho-Ser817 detected in our study is adjacent to the DM15 domain (877-915) in Chlamydomonas, a region in mammalian LARP1shown to be required for interaction with mTORC1 (Hong *et al*., 2017).

### Additional proteins with phosphosites altered by TORC1 inhibition

The majority of differential phosphosites we identified were not previously linked to TOR signaling, including in Chlamydomonas. These include sites on a translation-related protein (Cre17.g696250.t1.1) and RNA-binding proteins (Cre10.g441200.t1.2, Cre10.g466450.t1.1, Cre16.g659150.t1.1, Cre16.g662702.t1.1 Cre17.g729150.t1.2). One of the most down-modulated proteins annotated as CTC-interacting domain 4 (CID4, Cre01.g063997.t1.1), has been shown to have an important function in regulation of translation and mRNA stability in eukaryotes (Bravo *et al*., 2005; Jiménez-López *et al*., 2015). CID4 had 2 sites, Ser441 (FC=0.2_AZD8055_, FC=0.14_TORIN1_) and Ser439/Ser441/Ser446:NL (FC=0.03_AZD8055_, FC=0.05_TORIN1_) with a large decrease in phosphorylation upon inhibitor treatment. While little is known about the relationship between this protein and TORC1 signaling, the CTC domain, more recently referred to as the MLLE domain (Jiménez-López & Guzmán, 2014), is also found in evolutionarily conserved Poly (A)-binding proteins (PABPs). The large decrease in CID4 phosphorylation seen upon inhibition of the CrTORC1 pathway in our study implies a potential role for TORC1 mediated control of translation, similar to other well-known TOR substrates.

Another differential phosphosite of interest following TORC1 inhibition that was not previously linked to TOR regulation is a site on lycopene beta/epsilon cyclase protein (Cre04.g221550.t1.2--Thr800/Ser802:NL). This phosphosite is significantly increased upon Torin1 treatment (FC=4.02) and the total protein level remained constant upon rapamycin treatment (Supplementary Table S4, FC=0.88). Lycopene beta/epsilon cyclases are required for carotenoid biosynthesis, carrying out cyclation of lycopene to yield α- and β-carotenes (Cunningham *et al*., 1996; Cunningham & Gantt, 2001; Cordero *et al*., 2010) which have been shown to be high-value compounds participating in light harvesting and in the protection of the photosynthetic apparatus against photo-oxidation damage (Frank & Cogdell, 1996; Cunningham Jr & Gantt, 1998). Recently in rice, carotenoid content was shown to be significantly lower in an s6k1 mutant compared to wild-type (Sun *et al*., 2016) revealing a potential connection between the TOR pathway and carotenoid production. To further investigate the effect of TORC1 inhibition on carotenoid biosynthesis in Chlamydomonas based on our phosphoproteomic finding, carotenoid levels in AZD-, Torin1- and rapamycin-treated cells were assessed after eight hours of treatment with three biological and two technical replicates (Figure 6, Table 2). After eight hours of treatment, there was a significant increase in various carotenoids measured in TOR-inhibited samples including β-carotene, which is directly downstream of cyclase activity (Figure 6, Table 2). While the effects on carotenoid biosynthesis and secondary metabolism following TORC1 inhibition required eight hours to become detectable, this is the first evidence that carotenoid production is modulated by TOR signaling in algae. Additionally, altered cyclase protein levels are not likely responsible for this finding since previous studies showed no change in lycopene beta/epsilon cyclase protein level after up to 24 hours of nitrogen stress (Cunningham Jr & Gantt, 1998; Valledor *et al*., 2014), a condition that is metabolically similar to TOR inhibition (Perez-Perez *et al*., 2010; Roustan *et al*., 2017).

**Table 2:** Carotenoid content in WT Chlamydomonas after 8 hours of treatment with Rapamycin, Torin1, or AZD8055 compared to control.

Carotenoids Content (mg g<sup>-1</sup> DW)
|  | Control | 500nM Rap | 500nM Torin | 700nM Azd |
| --- | --- | --- | --- | --- |
| Neoxanthin | 0.64±0.01 | 1.12±0.01* | 1.25±0.04* | 1.65±0.00* |
| Violaxanthin | 0.50±0.00 | 0.57±0.00 | 0.93±0.07* | 1.02±0.00* |
| Anteraxanthin | 0.04±0.00 | 0.12±0.00* | 0.16±0.01* | 0.11±0.00* |
| Lutein | 1.60±0.03 | 2.56±0.02* | 3.42±0.14* | 3.29±0.00* |
| B-carotene | 1.82±0.03 | 1.80±0.03 | 2.09±0.02* | 3.02±0.02* |

**Figure 6:**
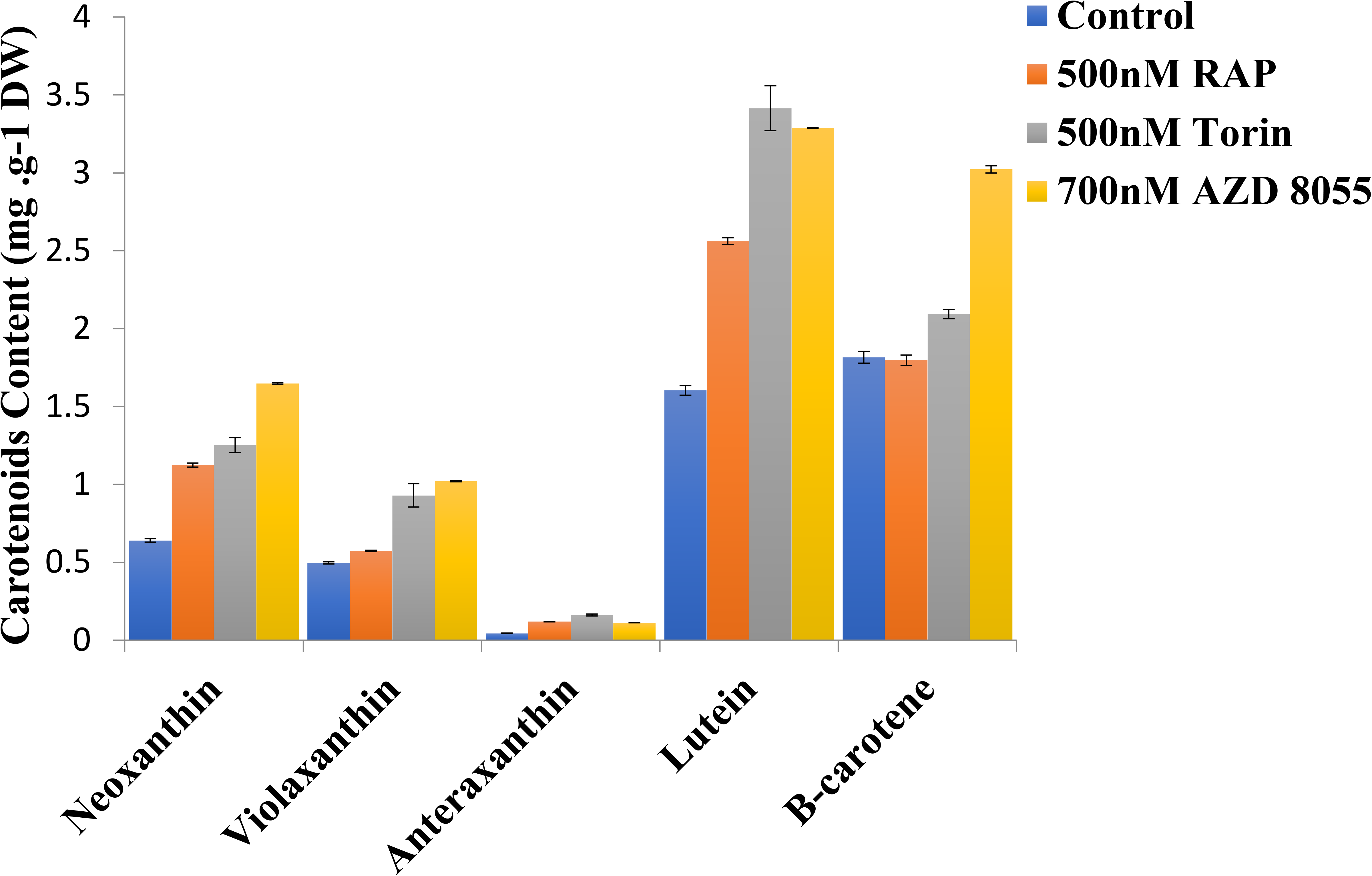
Bar chart of carotenoid content in WT Chlamydomonas after 8 hours of treatment with Rapamycin, Torin1, or AZD8055 compared to control

Numerous phosphosites from proteins without Phytozome database descriptions were also found to be down-regulated upon CrTORC1 inhibition, including some sites with large decreases (>five-fold). For all unannotated proteins, we searched for pfam, Panther, KOG, KEGG, KO, and GO pathway terms and domain conservation using Phytozome and NCBI annotations (Table S4). Numerous proteins had conserved domains including structural maintenance of chromosomes (Accession: cl25732), autophagy protein (Accession: cl27196), transmembrane proteins (Accession: cl24526), and small acidic protein (Accession: pfam15477). While the large changes upon chemical inhibition are potentially interesting, especially the five proteins containing sites with at least a five-fold decrease in phosphorylation (Cre03.g152150.t1.2, Cre06.g263250.t1.1, Cre11.g469150.t1.2, Cre05.g236650.t1.1, Cre13.g582800.t1.2), future targeted work would be required to infer biological significance to this observation. To aid in this, the fifty-eight modulated sites without Phytozome database annotation were also homology searched for best BLAST hit IDs in Volvox, Gonium, and Arabidopsis to find homologs among green lineage (Table S5) and Table S2 displays all of the experimentally derived sites modulated by AZD8055, Torin1, and/or rapamycin and will serve as a guide in follow-up studies.

In summary, we obtained a candidate list of phosphosites modulated following TORC1 inhibition. We achieved extensive coverage of the TOR-modulated phosphoproteome in Chlamydomonas using a quantitative label-free approach. Our approach was validated by the overlap of phosphosites altered using different TOR inhibitors and by our identification of Chlamydomonas homologs of TOR signaling-related proteins such as RPS6 and LARP1 that had decreased phosphorylation upon TORC1 inhibition. Follow-up experiments guided by our phosphoproteomic findings in lycopene beta/epsilon cyclase showed that carotenoid levels are affected by TORC1 inhibition, the first evidence that carotenoid production is under TOR control in algae. Conserved TOR substrate motifs were also identified such as RXXS/TP and RXXS/TP. Our study provides a new resource for investigating the phosphorylation networks governed by the TOR kinase pathway in Chlamydomonas.

## Acknowledgements

This research was supported by a National Science Foundation CAREER award (MCB-1552522) awarded to L.M.H.

## Author contributions

E.G.W., L.M.H., I.C.L., J.G.U., J.L.C. contributed to planning and experimental design. E.G.W., I. C.L., and Z.P. performed experiments. E.G.W., E.W.M. performed data analysis. E.G.W., L.M.H., J.G.U wrote the manuscript.

